# PathQC: Determining Molecular and Physical Integrity of Tissues from Histopathological Slides

**DOI:** 10.1101/2025.09.29.679347

**Authors:** Ranjit Kumar Sinha, Anamika Yadav, Sanju Sinha

**Affiliations:** Sanford Burnham Prebys Medical Discovery Institute, Center for Data Science, San Diego

## Abstract

Quantifying tissue molecular and physical integrity is essential for biobank development. However, current assessment methods either involve destructive testing that depletes valuable biospecimens or rely on manual evaluations, which are not scalable and lead to interindividual variation. To overcome these challenges, we present PathQC, a deep learning framework that directly predicts the tissue RNA Integrity Number (RIN) and the extent of autolysis from hematoxylin and eosin (H&E)-stained whole-slide images of normal tissue biopsies. PathQC first extracts morphological features from the slide using a recently developed digital pathology foundation model (UNI), followed by a supervised model that learns to predict RNA Integrity Number and autolysis scores from these morphological features. PathQC is trained on and applied to the Genotype-Tissue Expression (GTEx) cohort, which comprises 25,306 non-diseased post-mortem samples across 29 tissues from 970 donors, where paired ground truth RIN and autolysis scores were available. Here, PathQC predicted RIN with an average correlation of 0.47 and an autolysis score of 0.45, with notably high performance in Adrenal Gland tissue (R=0.82) for RIN and in Colon tissue (R=0.83) for autolysis. We provide a pan-tissue model for the prediction of RIN and autolysis score for a new slide from any tissue type (GITHUB). Overall, PathQC will enable scalable measurement of molecular and physical integrity from routine H&E images, thereby enhancing the quality of both biobank generation and its retrospective analysis.

## 1. Introduction

Traditional approaches to tissue quality assessment rely on molecular metrics, such as the RNA Integrity Number (RIN)[1], or manual histological evaluation of autolysis, both of which require either tissue consumption or extensive expert review [2]. The former is challenging when tissue samples are limited and adds sequencing cost, and the latter introduces inter-individual variations [3]. The digitization of pathology through whole slide imaging (WSI) technology has transformed how we capture and analyze tissue architecture. These high-resolution digital representations, typically scanned at 20× or 40× magnification, contain rich information about cellular morphology, tissue organization, and potentially, molecular integrity. However, current computational approaches in digital pathology primarily address technical image quality, detecting scanning artifacts [4], assessing focus quality [5], or normalizing staining variations [6] rather than investigating whether the visual patterns themselves might encode information about molecular integrity.

Recent advances, including ours, in computational pathology have revealed that histological images contain far more information than previously recognized [7, 8, 9, 10]. Foundation models trained on millions of pathology images, such as UNI [11], Virchow [12], and Prov-GigaPath[13], have demonstrated remarkable capabilities in extracting meaningful features from tissue morphology. Deep learning algorithms can now predict mutation status [14], gene expression profiles [15], microsatellite instability [16], telomere length [17], and treatment response directly from H&E-stained slides [18]. These discoveries challenge traditional boundaries between visual and molecular domains, suggesting a deeper connection between what we observe morphologically and what we measure molecularly.

Autolysis, the self-digestion of cells by their own enzymes, produces characteristic morphological changes, including nuclear pyknosis, cytoplasmic eosinophilia, and loss of cellular detail. Pathologists use these patterns to provide an autolysis score [19]. In the same vein, RNA degradation begins immediately after tissue devascularization, triggered by endogenous RNases released from lysosomes. This degradation follows predictable patterns: ribosomal RNA degrades first, followed by mRNA species with varying half-lives [20]. We hypothesize that a systematic analysis of tissue H&E images, paired with molecular RIN and autolysis score, can help us develop a machine learning model to determine it only from H&E images.

Towards this, we here present PathQC, a computational pathology quality framework to predict RIN number and autolysis score directly from H&E images. PathQC is trained and applied on the largest corpus of normal tissue data -Genotype-Tissue Expression (GTEx) cohort [21], with paired H&E and molecular quality metrics. The GTEx project, designed to study tissue-specific gene expression and regulation, provides an ideal foundation for our work with its 25,306 H&E-stained whole slide images spanning 29 distinct tissue types across approximately 970 individuals.

## 2. Results

### 2.1 Overview of PathQC Pipeline

The assessment of biological specimen quality, via RIN number and autolysis scores, requires a destructive sequencing assay and manual pathologist annotation. To overcome this, we here present PathQC (Figure 1A) to predict RNA Integrity Number (RIN) and autolysis scores directly from H&E-stained whole slide images (WSIs). The PathQC pipeline comprises four sequential steps (**Figure 1A**): 1) Slide Preprocessing, 2) Morphological feature extraction at the patch level, 3) Generating Whole-slide embedding, and 4) supervised model building to predict RIN and autolysis score based on morphological features. We provide a brief overview of these steps below:

**Figure. 1.**
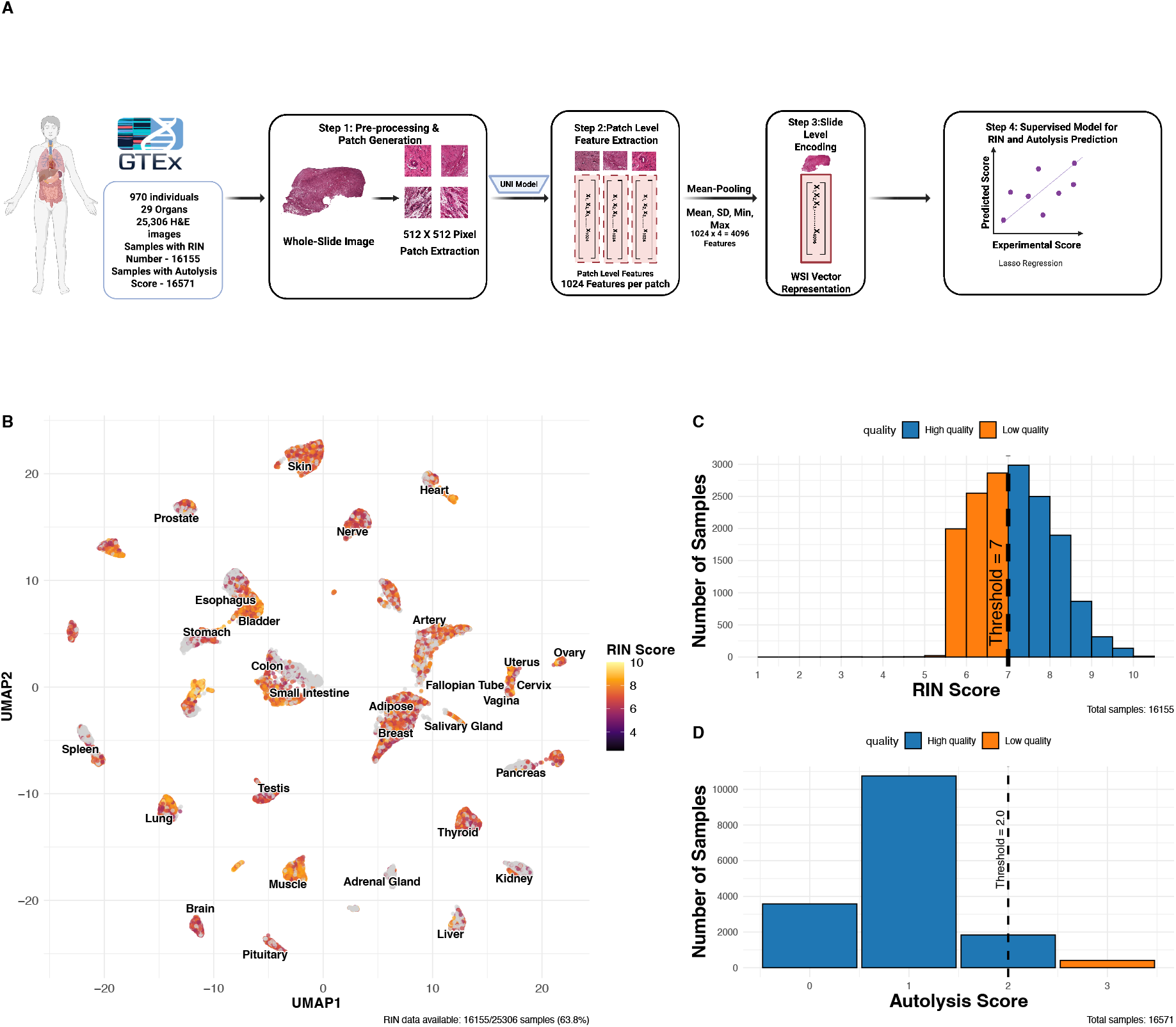
Overview of PathQC framework and tissue quality landscape. **A**. PathQC workflow showing preprocessing, feature extraction using UNI encoder, slide-level encoding, and supervised prediction. **B**. UMAP visualization with RIN score intensity overlay across 29 tissue types showing quality distribution patterns in morphological feature space. **C**. Distribution of RIN scores across the dataset showed that most samples had scores between 6-8. **D**. Distribution of autolysis scores showed most samples had scores of 0-1.

#### 1. Slide preprocessing

We begin with the preprocessing of whole slide images, removing background using Sobel edge detection and applying color normalization using Macenko’s method to mitigate batch effects. Images are segmented into 512×512-pixel patches, excluding those with insufficient tissue content (>50% background).

#### 2. Patch-Level Feature Extraction

For deep learning feature extraction, we process these normalized patches through the UNI encoder [10], a vision transformer architecture pre-trained on over 100 million tissue patches across multiple cancer types and normal tissues. This step yields 1,024-dimensional feature vectors for each patch, capturing essential visual patterns in each tissue region.

#### 3. Slide-Level Encoding

To bridge the resolution gap between patch-level features and slide-level quality metrics, we aggregate patches by taking their average, SD, max, and min, creating a 4096-length vector that creates a comprehensive slide-level feature vector that preserves essential visual information while enabling computational analysis.

#### 4. Developing a supervised model for RIN and Autolysis Scores

For quality prediction, we developed tissue-specific logistic regression models trained on GTEx RIN and autolysis scores. Their distribution is provided in **Figure 1C-D**. We compute our model’s performance in five-fold cross-validation, where the test set has never seen this data before.

We trained and applied PathQC using cross-validation, computing performance on test data never seen by the model. Our dataset comprised over 30 million patch images from 25,306 whole slide images across 29 tissue types from 970 donors, all with paired ground truth RIN and autolysis scores (**Figure 1C-D**). Our approach is based on the observation that in a UMAP based on morphological features, high vs. low RIN score samples are clustered separately (Figure 1B, Figure S1). This suggests that morphological features contain information to determine RIN scores, motivating our predictive modeling approach.

### 2.2 Landscape of Tissue Quality across different tissue types

Before building an H&E-based predictor, we next aim to understand how these RIN and autolysis scores vary in different tissues and their key confounding factors. To this end, we examined their variation across tissue types, demonstrating considerable tissue-specific variation, reflecting intrinsic differences in RNA integrity and degradation susceptibility **(Figure S6)**. We next examined technical confounders of these two metrics. (**Figure 2A, B**). For RIN scores, tissue type explained the largest portion of variance (15.2%), followed by the Hardy scale (circumstances of death) (11.6%), and the autolysis score (6.1%) (**Figure 2A**). Age and Gender had minimal impact, explaining only 3.3% and 0.6% of variance, respectively. For autolysis scores, tissue type explained an even larger variance proportion (27.7%), followed by RIN score (6.1%) and Hardy scale (4.3%) (**Figure 2B**). The analysis demonstrated that age had minimal influence on RIN scores across most tissue types, with tissue-specific variations observed in certain anatomical contexts. These results demonstrate the primary importance of tissue-specific characteristics in determining quality metrics, suggesting that quality assessment approaches should be calibrated to specific anatomical contexts rather than applied uniformly across tissues. Correlations between RIN and autolysis scores varied significantly across tissues **(Figure 2C)**, with the adrenal gland showing the strongest correlation (r = -0.59), indicating tissue-specific relationships between morphology and RNA integrity.

**Figure. 2.**
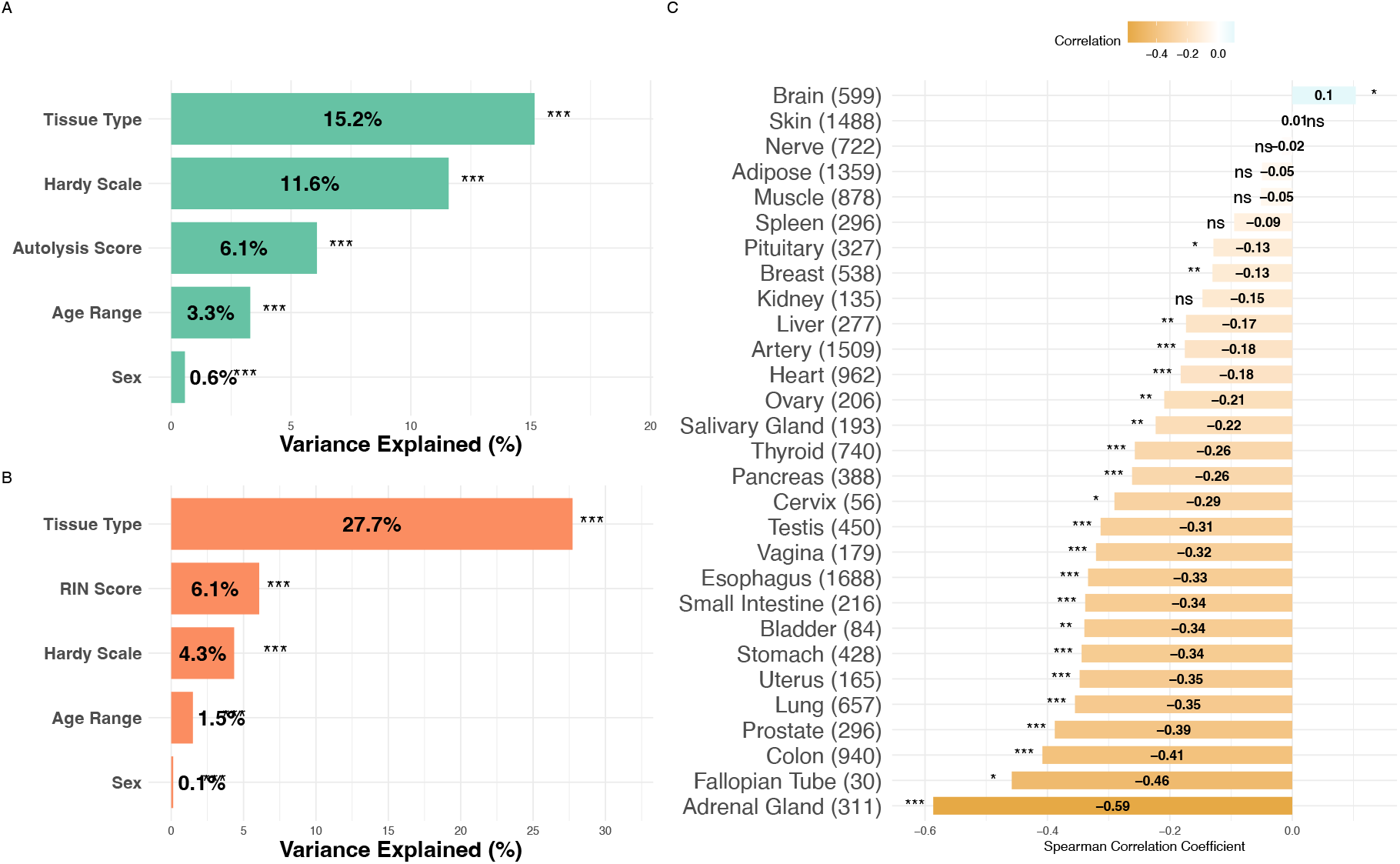
Factors affecting tissue quality in GTEx samples. **A**. For RIN scores, tissue type explained the largest portion of variance (15.2%), followed by the Hardy scale (circumstances of death) (11.6%), and autolysis score (6.1%). Age and Gender had minimal impact, explaining only 3.3% and 0.6% of variance, respectively. **B**. For autolysis scores, tissue type again explained the largest variance proportion (27.7%), followed by RIN Score (6.1%) and the Hardy scale (circumstances of death) (4.3%). **C**. Correlations between RIN and morphological features varied significantly across tissues, with the adrenal gland showing the strongest correlation with age (r = -0.59), indicating tissue-specific relationships between morphology and RNA integrity.

### 2.3 H&E morphology can determine RIN and autolysis score in 29 tissue types

We next built a H&E morphology-based predictor (fourth step of PathQC), a lasso regression model, using the above-noted sequencing-based RIN number and an expert pathologist’s annotation-based autolysis score in ∼25,306 GTEx biopsies. Our model was tested in five-fold cross-validation, where, across five iterations, the model’s performance was evaluated on 20% of the data that had never been seen by the model. PathQC predicted RIN score across 29 tissues with an average correlation of 0.47 and autolysis with an average correlation of 0.45. The prediction performance varies substantially across different tissue architectures, with RIN prediction ranging from 0.81 (Adrenal Gland), followed by esophagus (R = 0.77, RMSE = 0.75), liver (R = 0.72, RMSE = 0.67), heart (R = 0.72, RMSE = 0.69), to 0.17 (Nerve) (**Figure 3A, Figure S1**). Scatter plots for top-performing tissues (**Figure 3C**) illustrate the relationship between predicted and actual RIN scores. Similarly, autolysis performance ranged from 0.83 (Colon) to 0.13 (Nerve) (**Figure 3B**), and the predicted vs. true autolysis scores are provided for a few tissues in (**Figure 3D**). Our model’s cross-validation stability over many splits was tested in Notes S1 (Figure S7). We also observed a moderate positive correlation (r = 0.35) between RIN and autolysis prediction performance (**Figure S8A**), suggesting that quality metrics share some morphological manifestations.

**Figure. 3.**
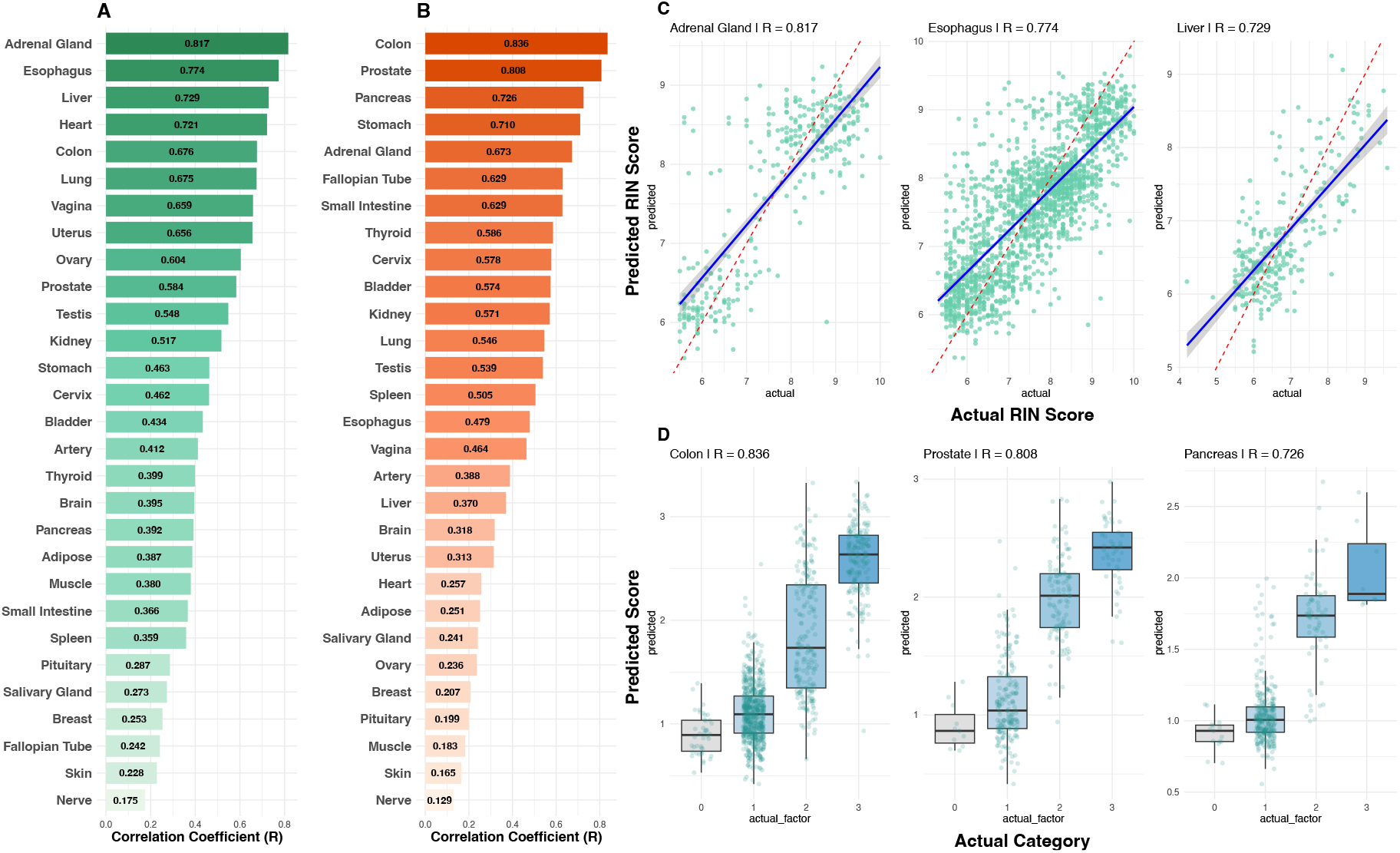
**A**. RIN prediction performance varied substantially across tissues, ranging from R=0.81 (adrenal gland) to R=0.17 (nerve), demonstrating that tissue biology rather than sample size drives predictability.**B**. Autolysis prediction performance ranged from R=0.83 (colon) to R=0.13 (nerve). Performance patterns revealed tissues with strong prediction for both metrics (colon, adrenal gland), metric-specific performance (esophagus for RIN; prostate for autolysis), and poor prediction for both metrics (nerve, skin). **C**. Scatter plots for top-performing RIN models showed clear correlations between predicted and actual values. **D**. Box plots for top-performing autolysis models demonstrated accurate classification into quality categories with minimal overlap between degradation levels.

### 2.4 Pan-Tissue model to jointly predict both integrity measures

Probing important features shared across tissue types, we identified the 20 most important features across all tissues for each integrity task (methods, Figure 4A-B). Hierarchical clustering revealed that morphologically similar tissues utilize similar feature sets. Further, 12 out of 20 top features were shared between RIN and autolysis models (Figure 4C). This prompted us to develop a pan-tissue model that can take any tissue type, H&E, and jointly predict both RIN and autolysis scores. This can be useful for an individual to use PathQC without relying on choosing tissue-specific models.

**Figure. 4.**
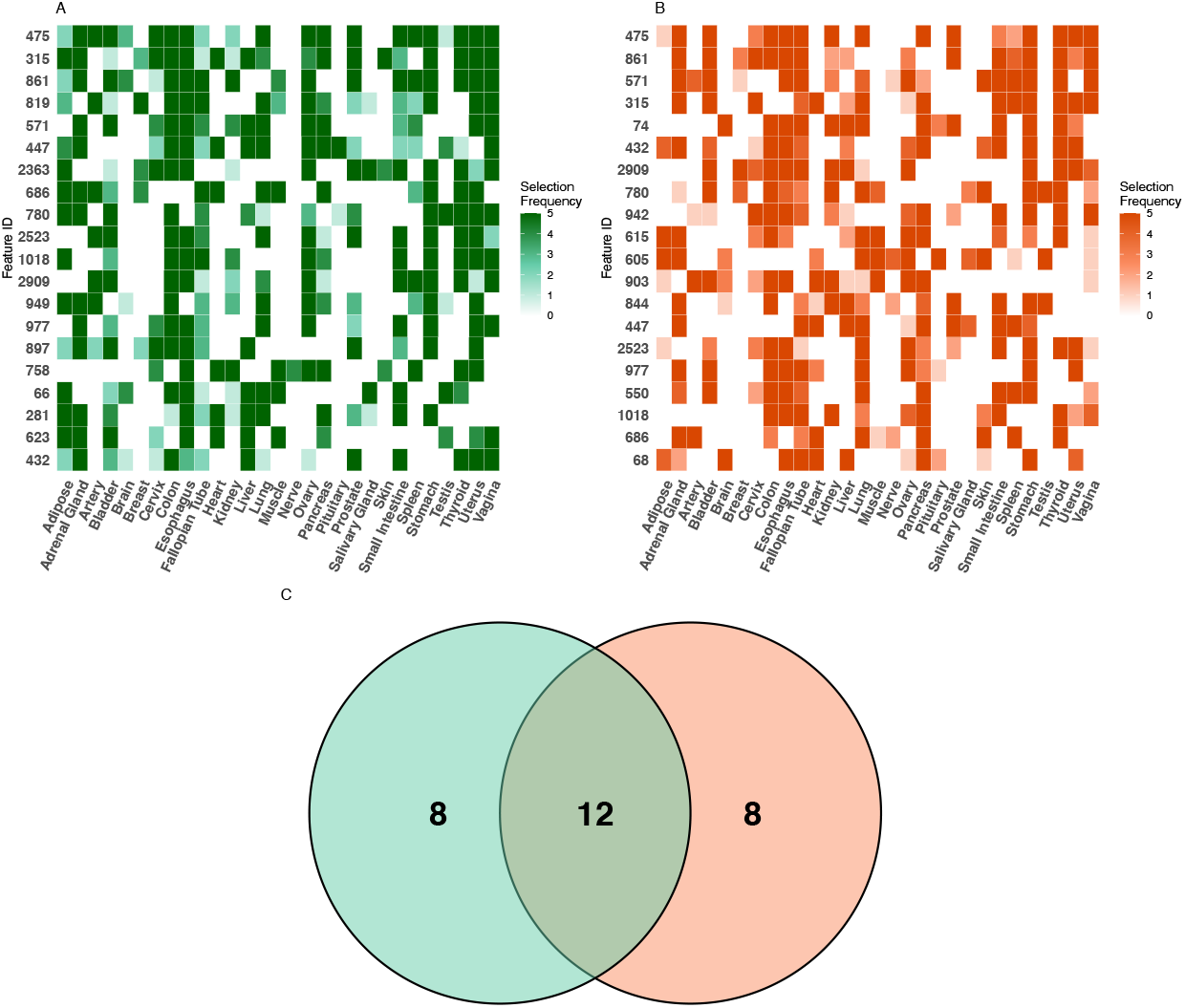
Feature importance analysis across tissue types. The feature importance heatmap illustrates the tissue-specificity of predictive features for **A**. RIN and **B**. Autolysis prediction. **C**. Comparing features important for both quality metrics, we identified 12 common features that showed high importance for both RIN and autolysis prediction across multiple tissues. These shared features suggest common morphological manifestations of degradation that affect both RNA integrity and general tissue preservation. Additionally, 8 features showed high specificity for RIN prediction, and another 8 features showed specificity for autolysis prediction, indicating that these quality aspects also manifest through distinct morphological changes.

To develop a pan-tissue model, we implemented a two-stage approach, where we first predict the tissue type and then accordingly select a tissue-specific model for prediction (Figure 5A). To this end, for our first step of tissue-type prediction, we developed a multinomial Lasso regression, where the top 500 most informative features are used (F-score). Our model predicted the right tissue type with 99.2% overall accuracy in 5-fold stratified CV (macro-averaged AUC >99.9%). Pairwise AUC analysis demonstrated exceptional tissue discrimination, with values ranging from 0.97 to 1.00 across all tissue pairs, indicating near-perfect separability based on morphology.

**Figure. 5.**
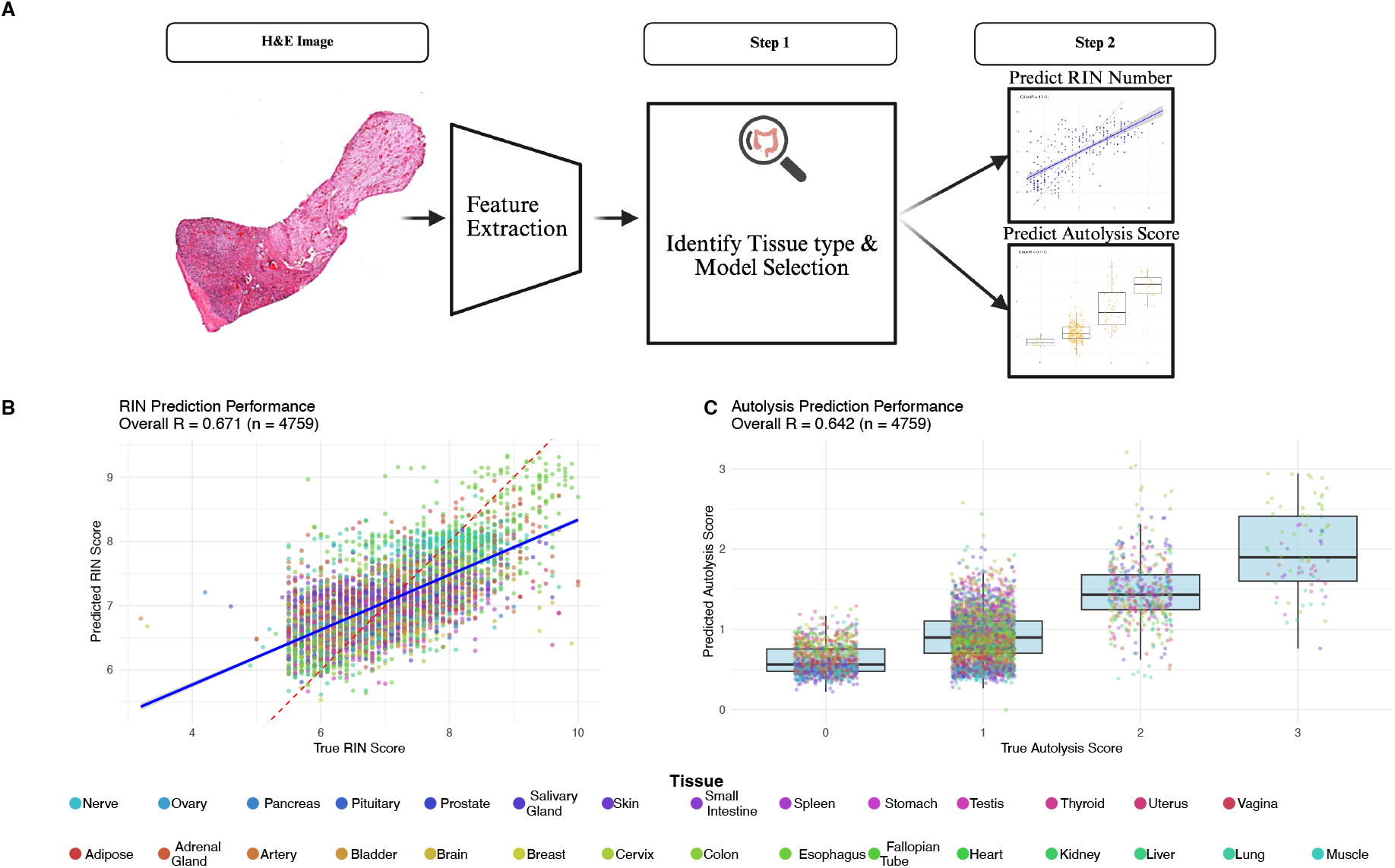
Pan-tissue model performance and workflow. **A**. Pan-tissue PathQC framework implementing a two-stage approach: tissue type classification followed by tissue-specific quality prediction, achieving 99.2% tissue classification accuracy with samples routed to appropriate tissue-specific models based on predicted tissue type. **B**. Pan-tissue RIN prediction achieved R = 0.67 (RMSE = 0.70) across 99.6% of test samples, with several tissues demonstrating consistent performance across both training and test sets. **C**. Pan-tissue autolysis prediction observed 70.1% prediction accuracy and R = 0.64 correlation, with tissues like Colon and Prostate showing robust morphological patterns for quality prediction across quality metrics.

Following tissue classification, samples were routed to appropriate tissue-specific models. The pan-tissue RIN prediction achieved R = 0.67 (RMSE = 0.70) across 99.6% of test samples (**Figure 5B**). For autolysis prediction, we observed 70.1% prediction accuracy and R = 0.64 correlation (**Figure 5C**). Several tissues demonstrated consistent performance across both training and test sets, including Colon and Prostate, suggesting robust morphological patterns for quality prediction across quality metrics **(Figure 5B-C)**. Comprehensive training and test performance metrics across all tissues within the pan-tissue framework, including correlation coefficients, sample sizes, and RMSE values for both RIN and autolysis predictions, are provided in **(Table S1)**.

## 3. Discussion

Building on the recent advancements in digital pathology foundation model to extract morphology features, we provide the first computational framework to predict RIN and autolysis score, tissue integrity measures, from H&E images. For biobanking workflows, PathQC adoption will enable rapid quality screening of large collections, preserve limited material for research rather than quality control, and facilitate retrospective assessment of historical collections where molecular quality data may be unavailable. Especially, a pan-tissue single model is applicable for most tissues and thus will help in adoption. Considering only 4% of total H&E samples in the U.S. are currently scanned, and this will only increase, adoption of digital pathology models and their applications will increase. PathQC serves as a basis for automated quality control using scanned images.

Despite its promising results, PathQC faces three key limitations: 1) The prediction performance varies substantially across tissue types, with some tissues showing excellent prediction while others remain challenging. 2) PathQC predicts bulk-tissue quality metrics rather than spatial quality variation within samples, where tissue degradation often occurs heterogeneously. 3) The interpretability of deep learning features remains challenging despite our feature importance analysis (**Figure 4A-B, S5**). The most promising direction for PathQC beyond addressing these three limitations with either technical development or additional data modalities is development of models that can conceptually take a step beyond predicting tissue damage and actually repair these tissues. Analogous to image-enhancing models in computer vision, enhancing damaged photos, one can learn from the large corpus of high-quality images and degrade them artificially to create paired labels. This conceptual breakthrough can help biobanks overcome tissue damage during preparation or scanning in H&E images at least. Broader validation studies across different sample preparation protocols and imaging platforms will be essential for establishing PathQC’s generalizability and clinical utility, paving the way for wider adoption in biobanking and clinical workflows.

## 4. Methods

### 4.1 Acquiring GTEx Dataset and Quality Metrics

Genotype-Tissue Expression (GTEx) v10 was downloaded from https://gtexportal.org/home/API. Each digitized at high resolution (20x magnification, ∼0.5 microns per pixel), these H&E-stained slides represent one of the largest collections of paired histological-molecular data available. For paired data, autolysis scores were assigned by expert pathologists using a 4-point ordinal scale: 0 (none), 1 (slight), 2 (moderate), and 3 (severe), based on digitally scanned whole slide images created using Aperio Scanscope software (Leica Biosystems). RIN scores were determined using the Agilent 2100 Bioanalyzer system (Santa Clara, California) to quantify RNA integrity.

### 4.2 PathQC Pipeline

PathQC comprises following four sequential steps, where the first three generating slide-level encoders were originally designed in Yadav, Alavarez et al. (2025) [16]. However, we provide their brief description here for completeness:

#### 1) Image preprocessing

We first prepared the WSIs by segmenting them into 512×512-pixel patches after background removal using Sobel edge detection. RGB color normalization using Macenko’s method.

#### 2) Patch-Level feature Extraction

A digital foundation model UNI, a pre-trained transformer-based encoder, was used to extract 1,024-dimensional feature vectors capturing morphology patterns in each patch.

#### 3) Generating Slide-level Embedding

Considering our quality metrics were at slide level, we generated slide-level embedding to train a predictive model for them. To this end, we aggregated patch features using statistical measures (mean, standard deviation, minimum, and maximum), creating a 4,096-dimensional slide-level feature vector.

#### 4) Supervised Training

We next aim to predict molecular quality metrics from above-created slide-level embedding. To this end, we developed tissue-specific Lasso regression models for both RIN and autolysis scores. Lasso regression was chosen for its ability to perform feature selection while building the predictive model, effectively handling the high-dimensional feature space while producing sparse, interpretable models. We z-score each feature and removed zero variance ones. The model was trained in a 5-fold cross-validation framework. Within each fold, top 5% features most correlated with target were selected, followed by a Lasso regression. Performance was evaluated using the correlation coefficient (R).

### 4.3 Feature Importance and Sharing Analysis

To analyze feature sharing patterns across tissues, we extracted feature selection frequencies from all tissue-specific models. For each target (RIN and autolysis), we aggregated feature selection counts across all tissues and identified the 20 most frequently selected features globally.

Feature usage patterns were visualized using heatmaps showing selection frequency by tissue type. To identify relationships between tissue types based on feature usage, we performed hierarchical clustering on both features (rows) and tissues (columns) using Euclidean distance and complete linkage. Feature overlap between RIN and autolysis models was quantified by comparing the top 20 feature sets from each prediction task.

### 4.3 Model Stability and Cross-Validation Analysis

To evaluate model consistency and reliability across folds, we conducted stability analysis across cross-validation folds by calculating the coefficient of variation (CV) of correlation coefficients for each tissue-specific model: **CV = σ/μ**,

where σ is the standard deviation and μ is the mean of correlation coefficients across folds. Models with CV values below 0.2 were classified as stable, while those exceeding 0.5 were considered unstable.

### 4.4 Pan-Tissue Model Development

To enable quality prediction across diverse tissue types, we developed a hierarchical pan-tissue framework consisting of tissue classification followed by tissue-specific quality prediction. Dataset Preparation: We combined samples from all 29 tissue types with both RIN and autolysis scores (n = 14,539), applying a 70:30 train-test split stratified by tissue type and RIN quartiles.

#### 1. Tissue Classification

ANOVA F-score selection identified the top 500 discriminative features across tissue types. We trained a multinomial Lasso regression model using stratified 5-fold cross-validation with L1 regularization (α = 1).

#### 2. Quality Prediction

Samples were routed to tissue-specific models based on predicted tissue type and a confidence threshold (0.6). Separate Lasso regression models for RIN and autolysis were trained for each tissue using all 4,096 features with stratified cross-validation by RIN quartiles and autolysis categories, respectively.

Performance Evaluation: Tissue classification was assessed using accuracy, macro F1-score, and macro-AUC. Quality prediction employed correlation coefficients for RIN and classification accuracy, along with correlation for autolysis. Coverage was calculated as the proportion exceeding the confidence threshold.

## 5. Author Contributions

SS conceptualized the framework of predicting RIN Numbers and Autolysis Scores from H&E. RKS developed the computational framework, performed data analysis and model validation, and contributed to manuscript writing. AY performed feature extraction and data preprocessing. SS supervised the project and mentored manuscript writing. All authors read and approved the final manuscript.

## 6. Data Availability Statement

The data used in this study were obtained from the Genotype-Tissue Expression (GTEx) Project (accession: phs000424.v10.p2, Project titled #39103). The H&E histological images are publicly available and were accessed through the GTEx Portal v10 release (https://gtexportal.org/home/). Autolysis Score and RIN score were also obtained from GTEx Portal (https://www.ncbi.nlm.nih.gov/gap/). Individuals age were extracted after appropriate institutional approval and data access agreement (General Research Use: phs000424.v10.p2.c1). All processed data for figure generation is provided in supplementary tables.

## 7. Code Availability

The underlying code for this study is available on GitHub and can be accessed via https://github.com/Cranjit9/PathQC

## 8. Competing Interest Statement

The authors have declared no competing interests.

