## Supplementary material for "PathQC: Determining Molecular and Physical Integrity of Tissues from Histopathological Slides": Supplemantary Text and figure

**Supplementary Notes, Figures**

**Supplementary Notes**

1. Quality distributions across tissues reflect tissue-specific biological characteristics: Quality score distributions across major tissue types revealed tissue-specific patterns in both RNA integrity and degradation susceptibility (**Figure S6**). RIN score distributions showed that tissues like esophagus, artery, and skin demonstrated consistently high scores with narrow distributions, while prostate, kidney, and liver revealed broader distributions with lower median values (**Figure S6A**). These varying distribution shapes reflect tissue-specific patterns in RNA integrity, with some tissues showing narrow, peaked distributions indicating consistent preservation, while others exhibit broader ranges suggesting greater variability in RNA quality. Autolysis score distributions demonstrated that most tissues show predominantly minimal degradation (scores 0-1), with varying proportions of samples demonstrating moderate to severe autolysis (**Figure S6B**). Notable patterns included tissues like liver, kidney, and prostate showing broader autolysis ranges, suggesting greater susceptibility to post-mortem degradation.

2. *Testing cross-validation stability:* We next tested our model's cross-validation stability, demonstrating that our tissue-specific models achieved consistent performance across data splits (**Figure S7**). The coefficient of variation for R values across folds averaged 0.10 for high-performing tissues and 0.21 for moderate performers, indicating reasonable stability given the challenging nature of the prediction task. Further, sample size analysis **(Figure S2)** revealed no significant correlation between training set size and model performance ($r = -0.13$, $p = 0.31$). For example, skin showed poor performance for both RIN ($R = 0.22$, $n = 1{,}492$) and autolysis ($R = 0.16$, $n = 1{,}502$) prediction despite having the largest training sample sizes, while fallopian tube demonstrated variable performance with poor RIN prediction ($R = 0.24$, $n = 30$) but high autolysis prediction ($R = 0.62$, $n = 30$) using the same small sample size. In contrast, colon achieved excellent performance for both metrics (RIN $R = 0.67$, $n = 940$; autolysis $R = 0.83$, $n = 1{,}057$), demonstrating that tissue biology, rather than sample size, drives predictability. We next tested the stability of top features across cross-validation. This first revealed that top-performing tissues relied on more stable feature sets than poor-performing tissues (**Figure S11**, 74.1% vs 57.5% mean consistency across folds). For example, tissues with the highest consistency included esophagus (85.8%), colon (85.1%), and heart (84%), while tissues with lower consistency included fallopian tube (40.8%), nerve (42.2%), and cervix (44.2%), suggesting varying reliability of morphological indicators across tissue types.

3. Important Features Analysis: Analyzing top features from both models, we note that mean-based features dominated both prediction tasks with 45 features each, while maximum, minimum, and standard deviation categories contributed fewer features (**Figure S5A**).

4. *Testing Modelling assumptions*: Residual plots for top-performing RIN and autolysis-models confirmed that residuals were randomly distributed with minimal systematic bias (**Figure S4A-B**). Low absolute residuals (**Figure S4C-D**) confirms that model performance is not driven by a few outliers.

Supplementary Figures:

Figure S1. UMAP visualization of top-performing tissue quality prediction models.
Figure S2: Sample size effects on model performance across tissue types.
Figure S3: Feature consistency and cross-tissue analysis
Figure S4. Model residual analysis and prediction accuracy validation
Figure S5. Feature importance analysis and morphological pattern characterization
Figure S6. Quality score distributions across major tissue types
Figure S7. Cross-validation stability and model robustness evaluation
Figure S8. Comparative model analysis and cross-metric relationships.
Figure S9. Model performance comparison across top-performing tissues
Figure S10. Comprehensive analysis of tissue morphological patterns and dataset characteristics
Figure S11. Model performance vs In-fold consistency
Figure S12. Pan-tissue model performance validation across test data

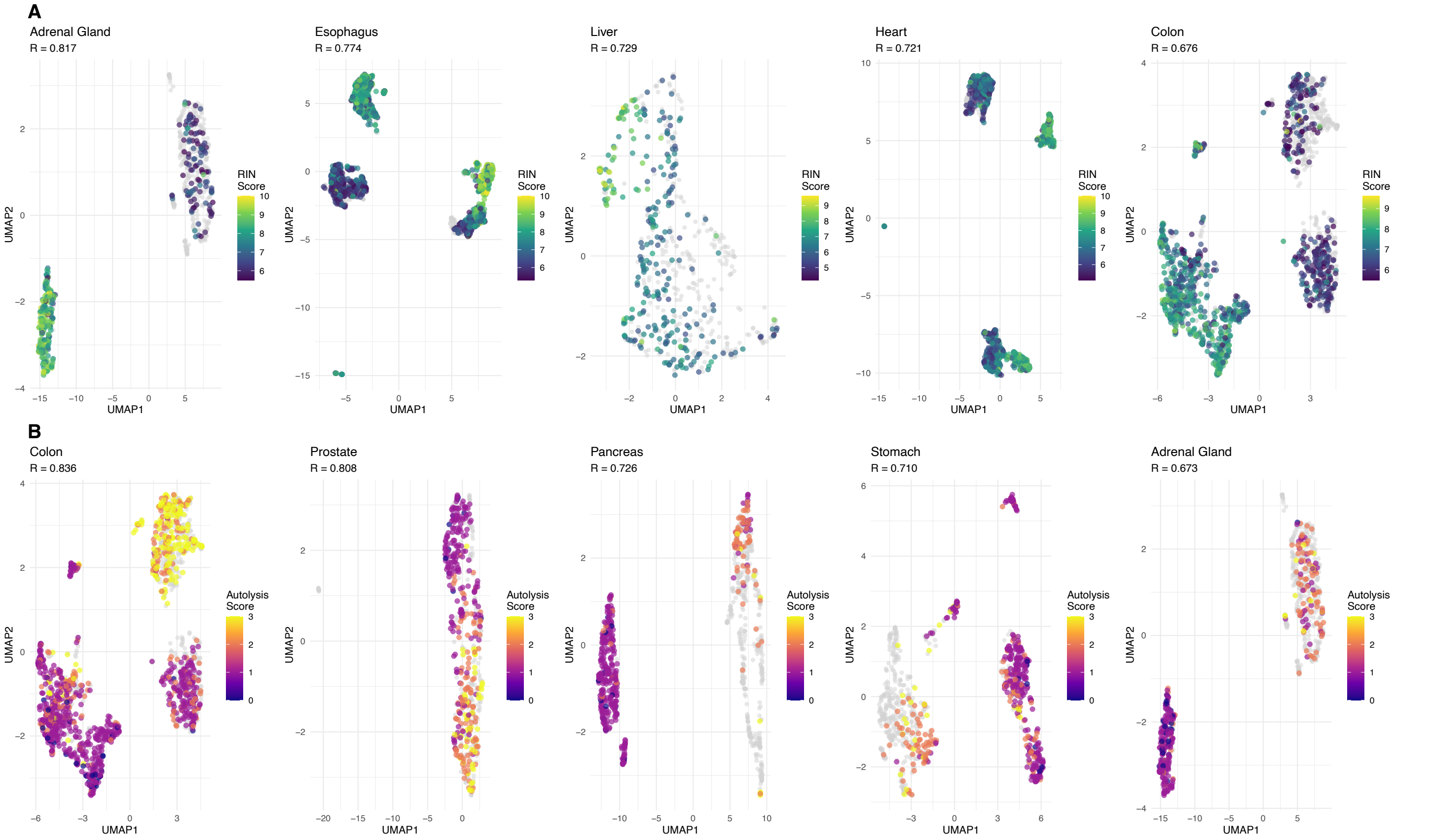


**Figure S1. UMAP visualization of top-performing tissue quality prediction models. A.** UMAP projections for top 5 RIN prediction models (Adrenal R=0.81, Esophagus R=0.77, Liver R=0.72, Heart R=0.72, Colon R=0.67). Samples with higher RIN scores cluster separately from degraded samples, validating morphological feature capture. **B.** UMAP projections for top 5 autolysis prediction models (Colon R=0.83, Prostate R=0.80, Pancreas R=0.72, Stomach R=0.71, Adrenal R=0.67). Clear spatial separation between minimal and severe degradation levels.

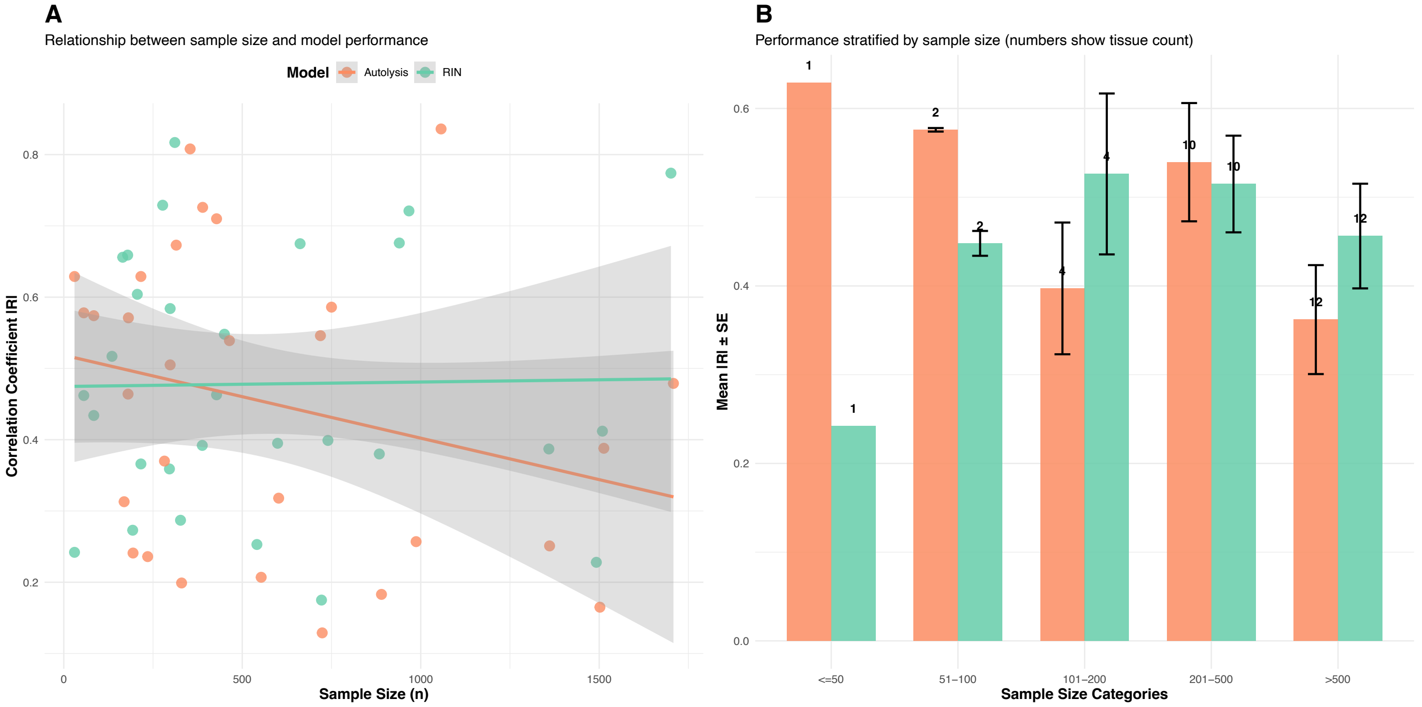

**Figure S2. Sample size effects on model performance across tissue types. A.** Scatter plot of training sample size vs model performance (|R|) for RIN and autolysis models across 29 tissues. Weak correlation suggests tissue biology drives predictability more than sample size. **B.** Performance stratified by sample size categories (≤50, 51-100, 101-200, 201-500, >500). Substantial overlap between categories confirms tissue-specific characteristics dominate over training data quantity.

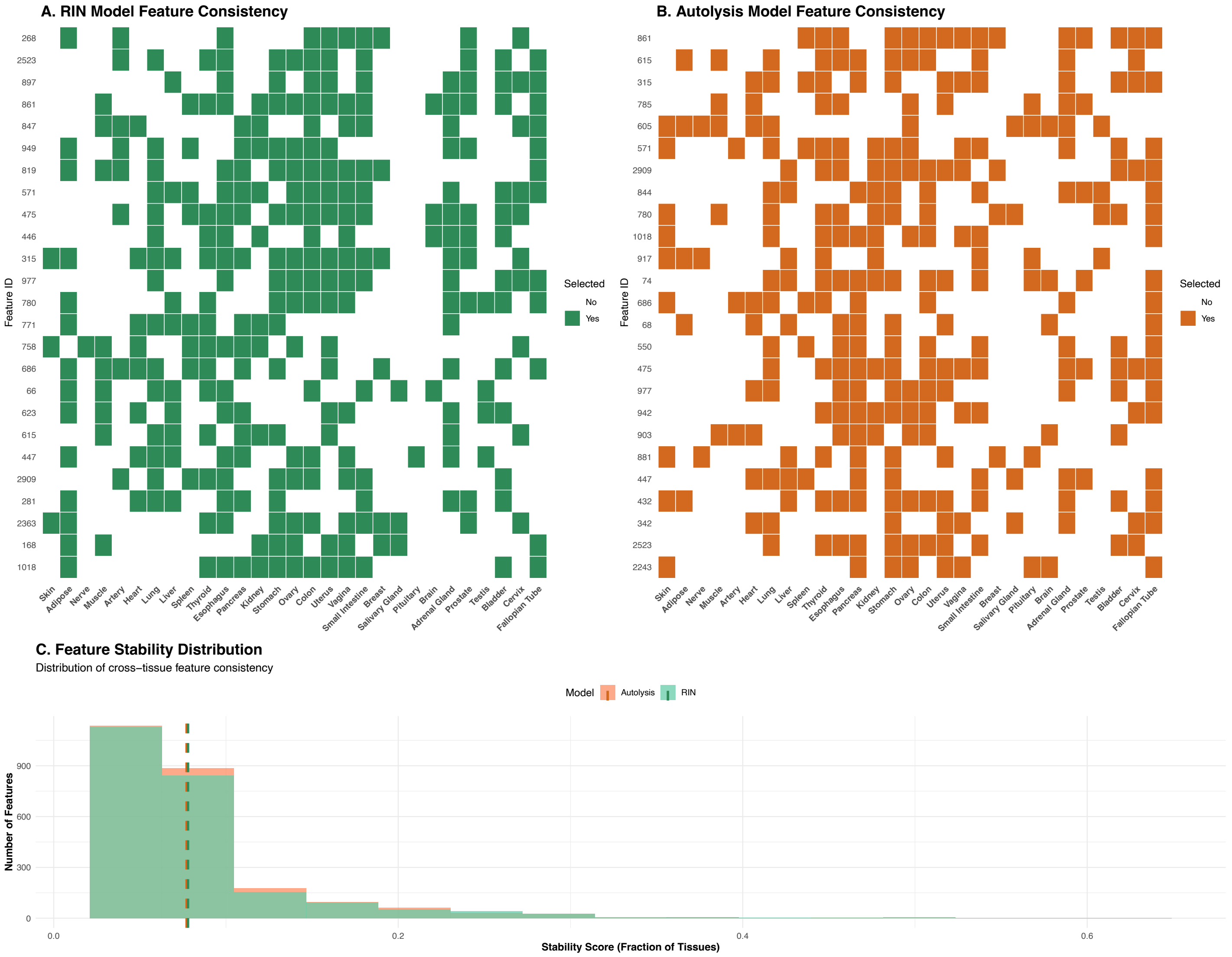


**Figure S3. Feature consistency and cross-tissue analysis. A.** Feature selection heatmap for RIN prediction across 29 tissues. Green squares show most features exhibit tissue-specific rather than universal importance. **B.** Feature selection heatmap for autolysis prediction with orange squares. Reveals distinct but partially overlapping patterns compared to RIN models. **C.** Distribution of feature stability scores comparing RIN and autolysis models. Most features show low cross-tissue consistency, validating tissue-specific modeling approach.

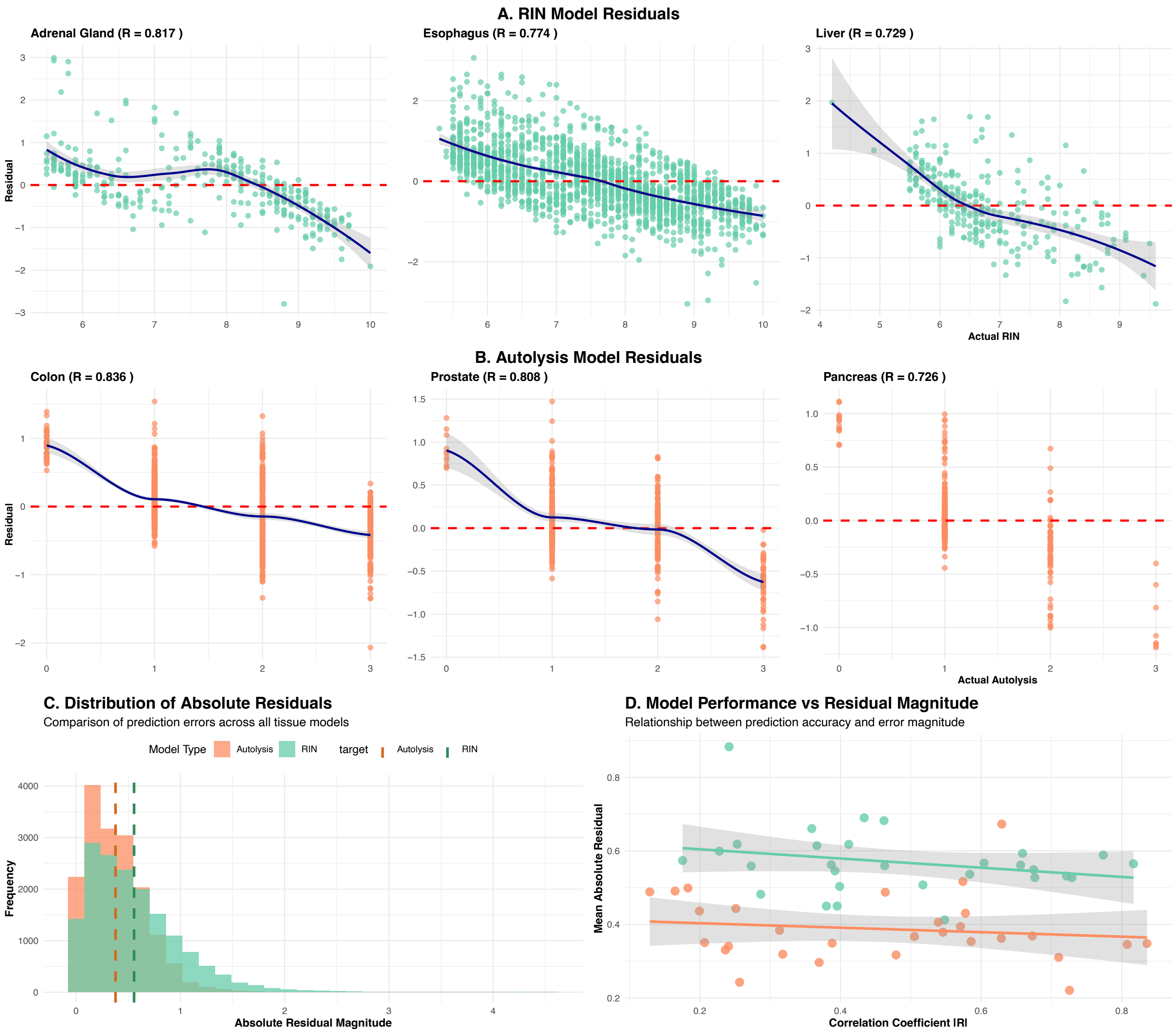


**Figure S4. Model residual analysis and prediction accuracy validation. A-B.** Residual plots for top-performing RIN and autolysis models. Random distribution around zero confirms well-calibrated predictions with minimal systematic bias. **C.** Histogram of absolute residual magnitudes for both model types. Most residuals concentrated in lower range (0-1) indicating accurate predictions. **D.** Scatter plot showing strong negative relationship between model performance and prediction error. Higher-performing models consistently exhibit smaller residuals.

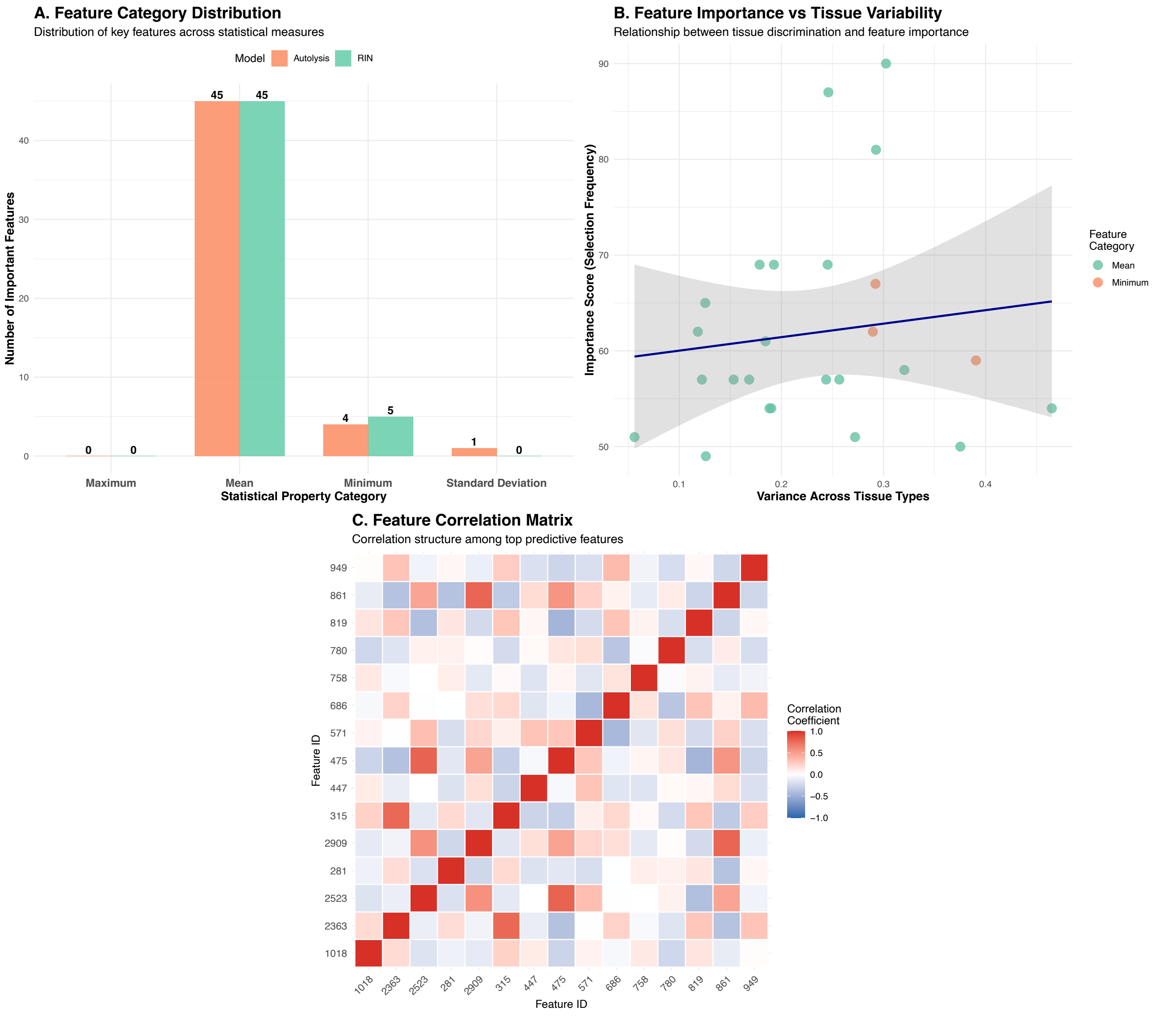


**Figure S5. Feature importance analysis and morphological pattern characterization. A.** Distribution of important features by statistical categories. Mean-based features dominate both tasks (45 each), indicating average tissue characteristics are most predictive. **B.** Scatter plot of feature importance vs tissue variability. Positive trend shows features varying across tissues are more frequently selected in models. **C.** Correlation matrix of top 15 predictive features. Blue-red scale reveals clusters of correlated morphological patterns.

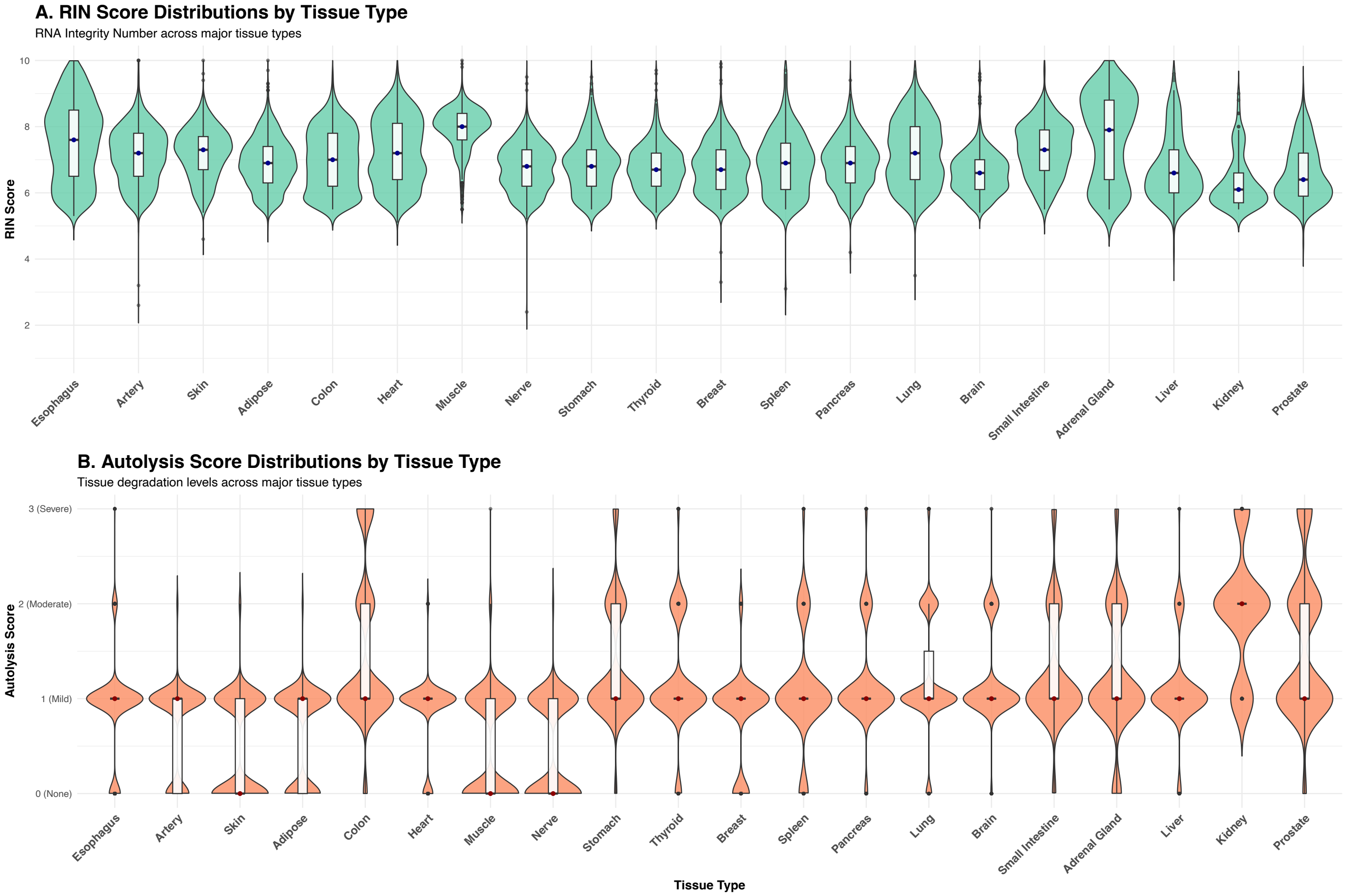


**Figure S6. Quality score distributions across major tissue types. A.** Violin plots of RIN scores across 20 tissues ordered by median values. The esophagus and artery show high scores with narrow distributions, while prostate and kidney show broader, lower distributions. **B.** Violin plots of Autolysis scores for same tissues. Most show minimal degradation (0-1), with liver, kidney, and prostate displaying broader ranges.

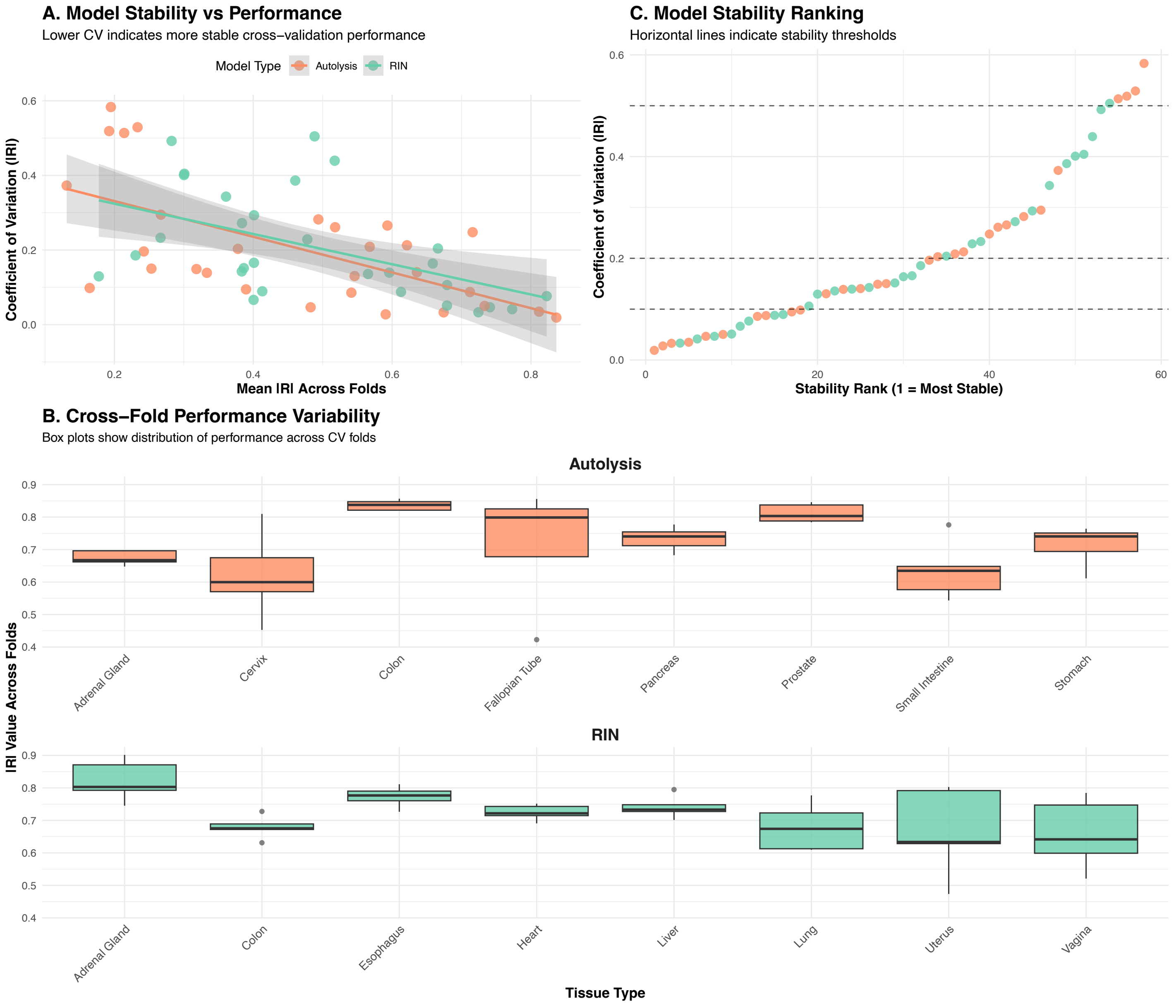


**Figure S7. Cross-validation stability and model robustness evaluation. A.** Scatter plot of stability (coefficient of variation) vs mean performance. Negative correlation confirms higher-performing models show greater robustness across data splits. **B.** Box plots of correlation coefficients across folds for selected tissues. Demonstrates consistent performance with narrow interquartile ranges for stable models. **C.** Model stability ranking by coefficient of variation. Approximately 70% of models achieve reasonable stability below 0.2 threshold.

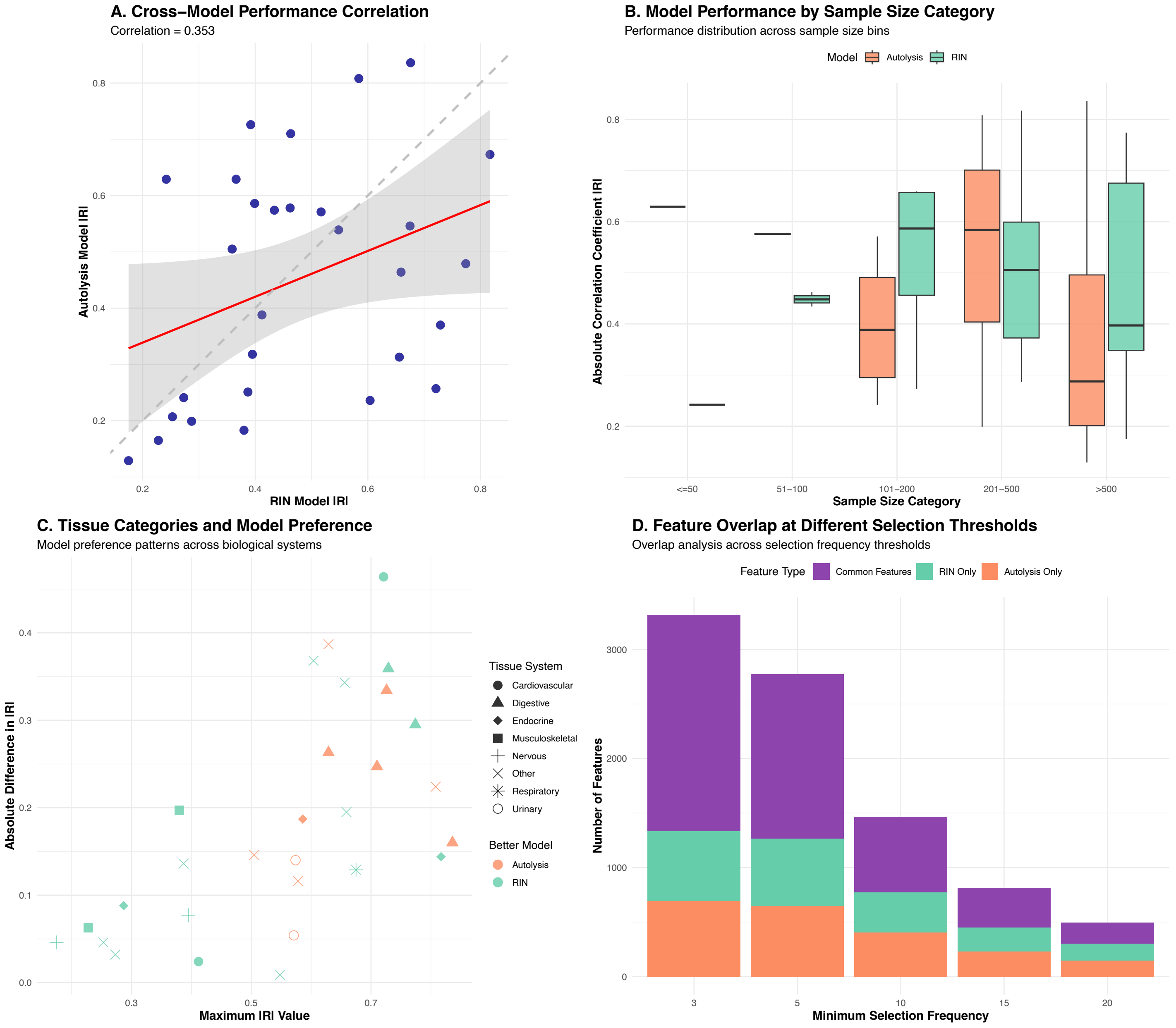


**Figure S8. Comparative model analysis and cross-metric relationships. A.** Scatter plot comparing RIN vs autolysis model performance across tissues. Moderate correlation (r=0.35) suggests shared but distinct morphological patterns. **B.** Box plots of performance by sample size categories. Substantial overlap reinforces that tissue biology is primary determinant over sample quantity. **C.** Analysis of tissue categories and model preferences by biological system. Points colored by system and shaped by preferred model type. **D.** Stacked bar chart of feature overlap at different selection thresholds. Shows model-specific features increase while common features decrease at higher thresholds.

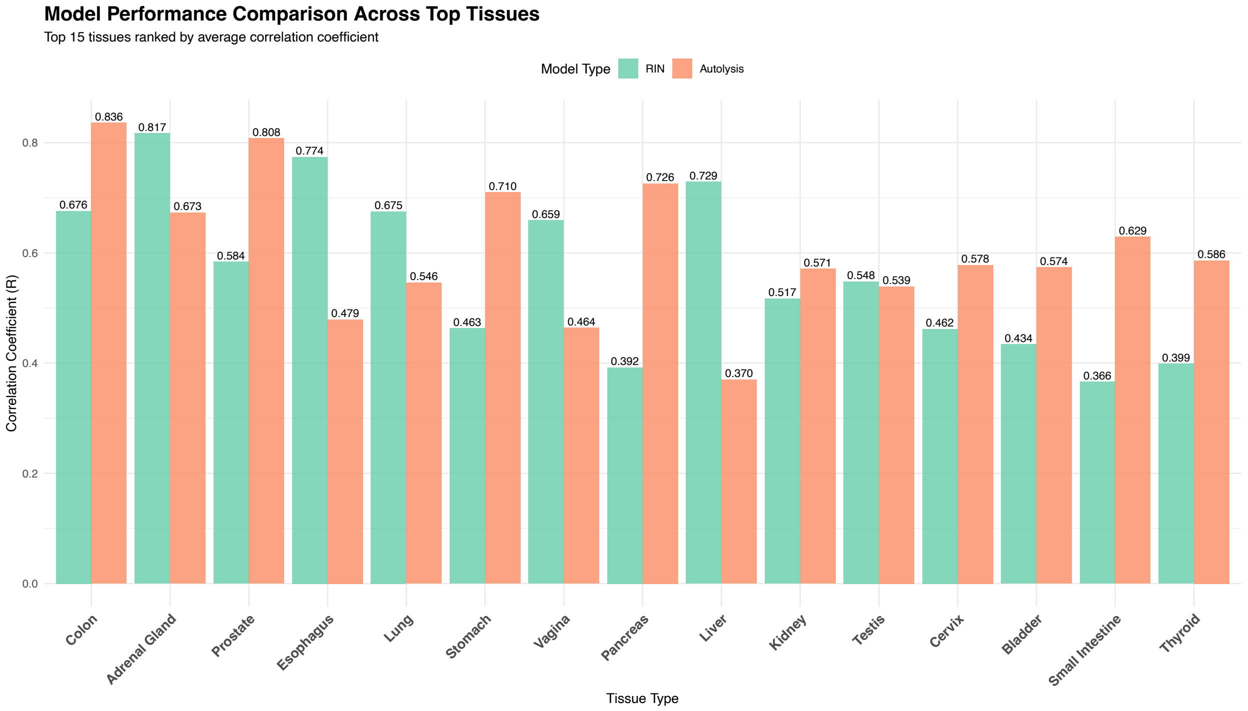


**Figure S9. Model performance comparison across top-performing tissues.** Bar chart comparing RIN and autolysis performance for top 15 tissues ranked by average correlation. Reveals three patterns: dual high performers (colon, adrenal), clear preferences (esophagus/liver for RIN; prostate/stomach for autolysis), and moderate dual performers.

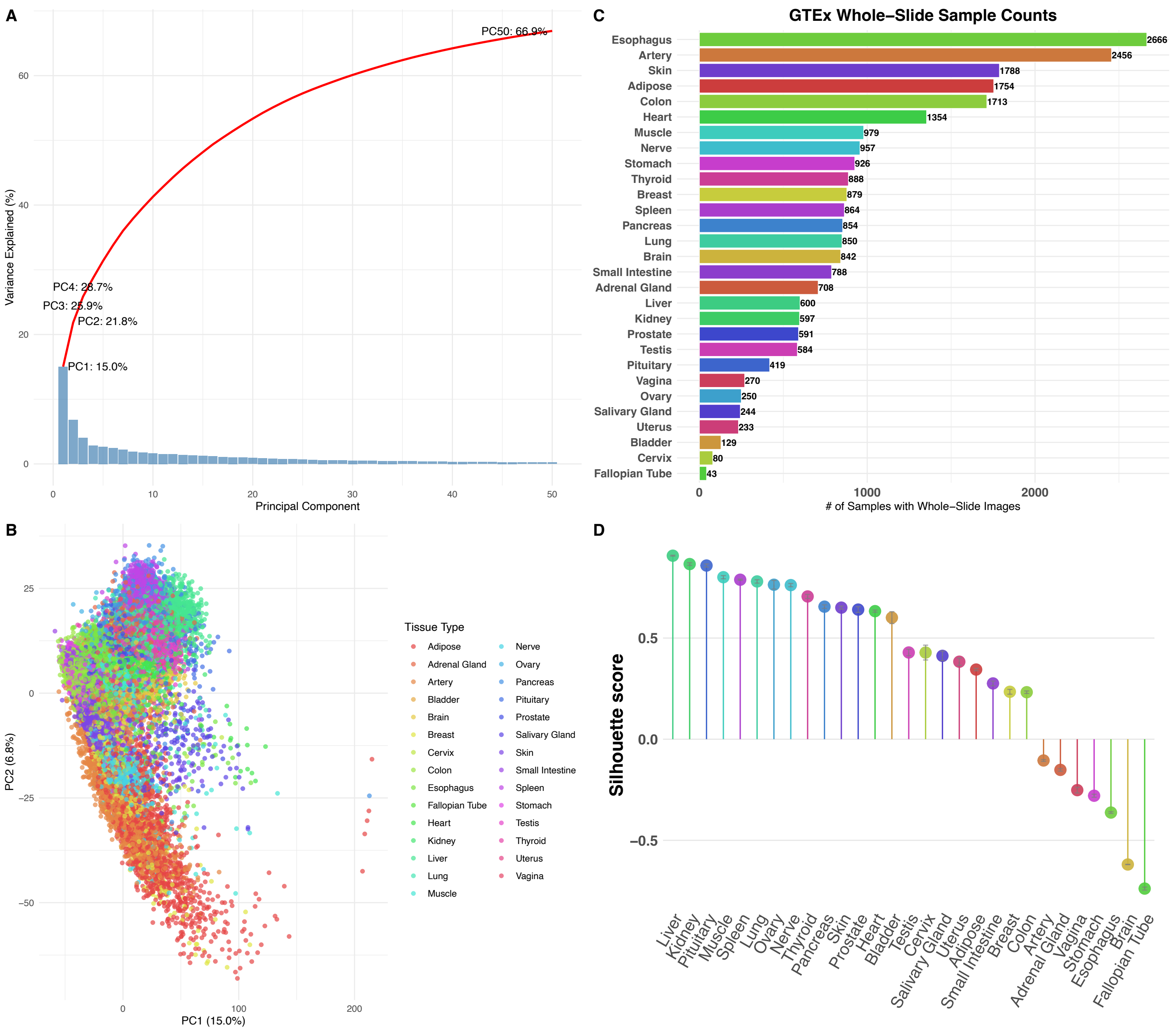


**Figure S10. Comprehensive analysis of tissue morphological patterns and dataset characteristics. A.** Scree plot of first 50 principal components. PC1 explains 15.0%, PC2 adds 6.8%, demonstrating high-dimensional complexity requiring many components. **B.** PC1 vs PC2 scatter of 25,306 GTEx samples colored by tissue type. Shows distinct clustering patterns reflecting tissue-specific morphological characteristics. **C.** Horizontal bar chart of sample counts across 29 tissue types. Esophagus largest (n=2,666), fallopian tube smallest (n=43). **D.** Silhouette analysis ranking tissues by clustering quality. Liver, kidney, pituitary show highest separation (>0.6), while brain, artery show overlap.

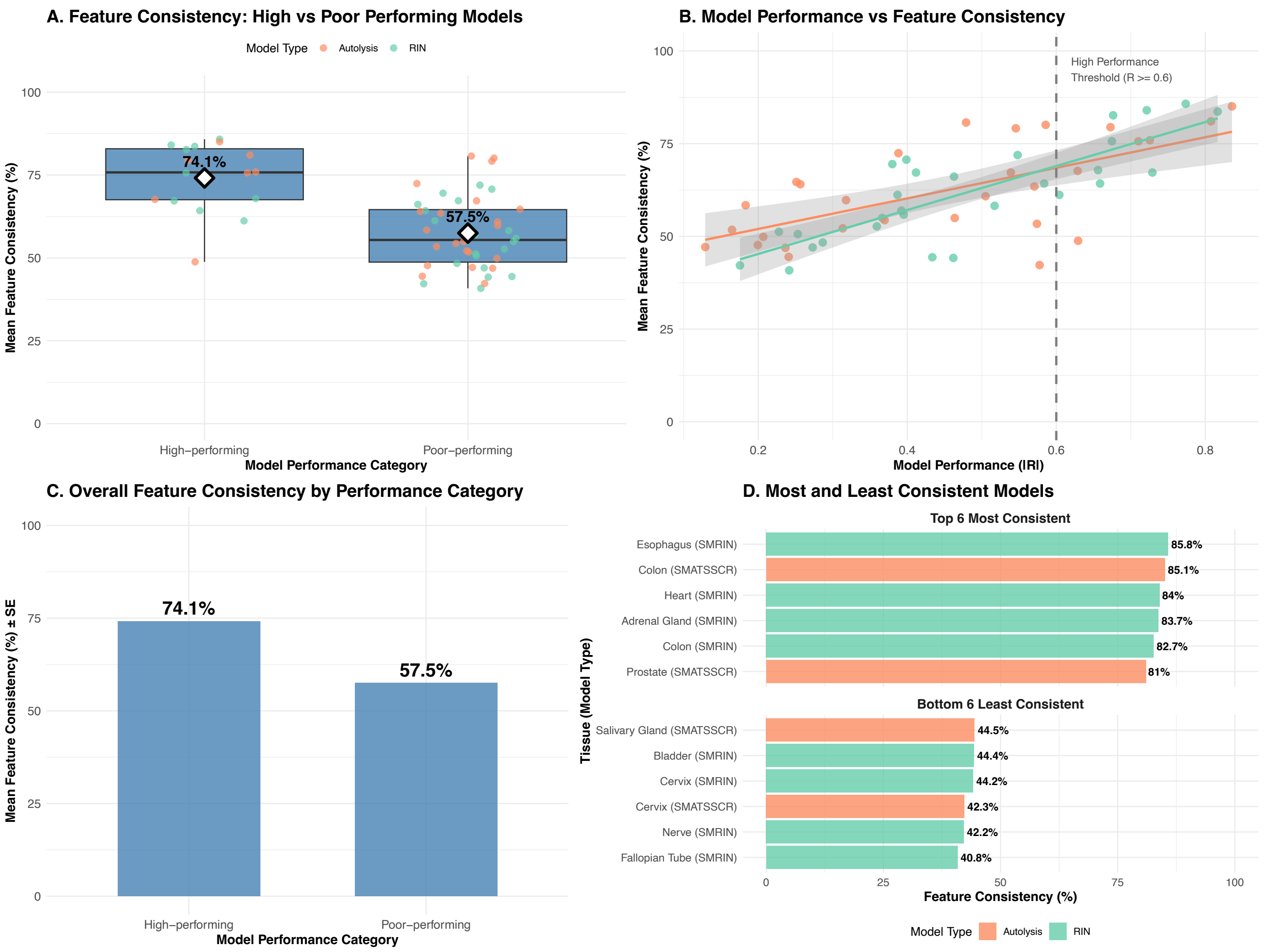


**Figure S11. Model performance vs In-fold consistency A.** Box plots comparing feature consistency between high-performing (|R|≥0.6) and poor-performing models. High performers show 74.1% vs 57.5% consistency, indicating stable feature selection drives robust prediction. **B.** Scatter plot of model performance vs mean feature consistency. Positive relationship confirms successful models achieve more stable patterns. **C.** Bar chart with error bars comparing overall consistency between performance categories. Confirms 74.1% vs 57.5% difference with statistical significance. **D.** Horizontal bar chart ranking tissue-model combinations by feature consistency. Esophagus (SMRIN) leads at 85.8%, while Bladder (SMRIN) shows lowest at 44.4%.

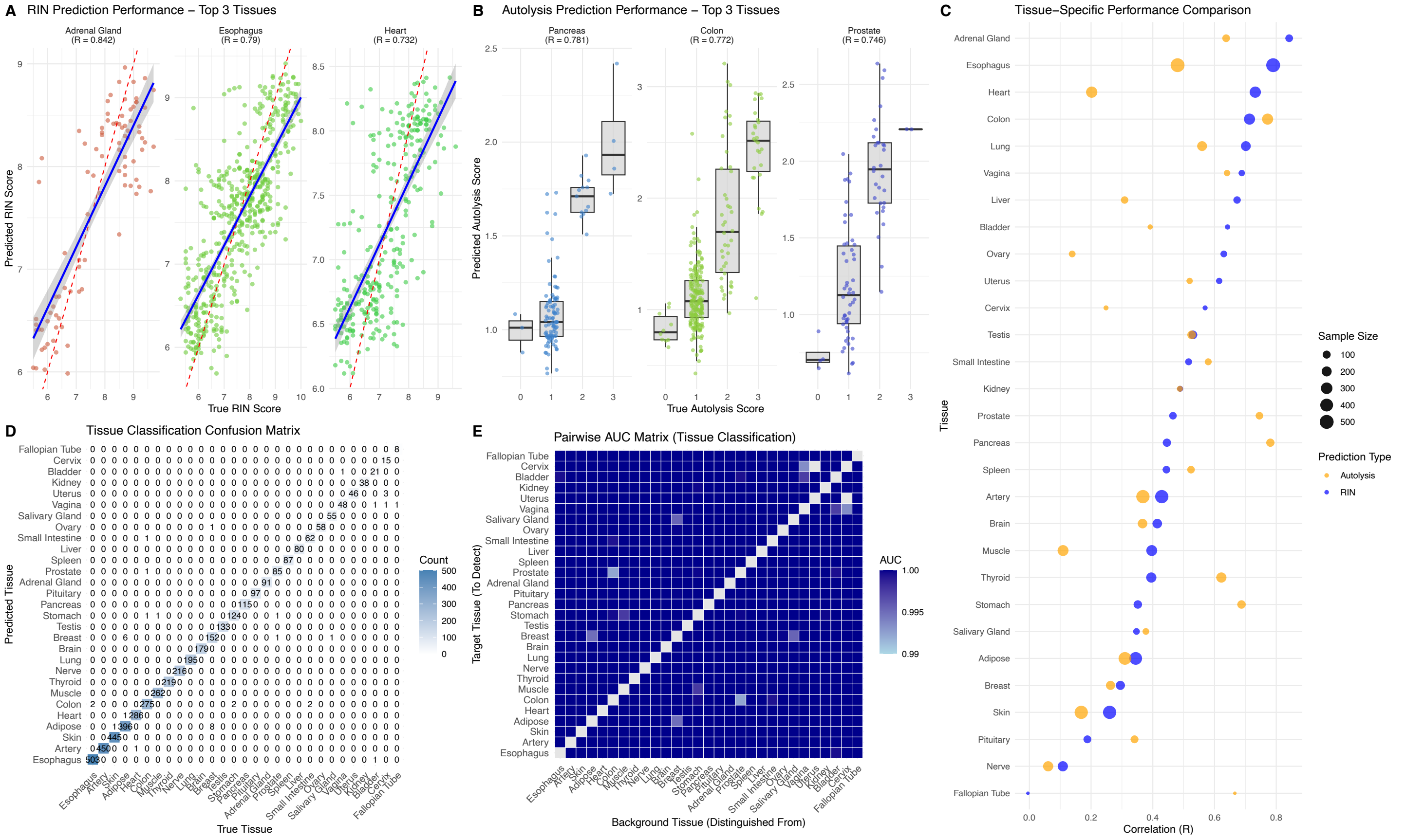


**Figure S12. Pan-tissue model performance validation across test data. A.** RIN prediction scatter plots for top 3 test performers: Adrenal Gland (R=0.842), Esophagus (R=0.790), Heart (R=0.733). **B.** Autolysis prediction boxplots for top 3 test performers: Pancreas (R=0.780), Colon (R=0.773), Prostate (R=0.746). **C.** Cross-tissue performance comparison showing RIN vs autolysis correlations. Point size indicates sample size; colors distinguish prediction types. **D.** Tissue classification confusion matrix demonstrating 99.2% accuracy across 29 tissue types. **E.** Pairwise AUC heatmap showing tissue discrimination performance ranging from 0.97-1.00.
