## Supplementary material for "PathQC: Determining Molecular and Physical Integrity of Tissues from Histopathological Slides": Supplemantary Table

Dear Drs. Jinman Kim, Adam Dunn & Lei Bi,

We are pleased to submit our manuscript entitled "PathQC: Determining Molecular and Physical Integrity of Tissues from Histopathological Slides" for consideration in your journal for “Multimodal AI for Digital Medicine” collection.

Tissue biobanks serve as foundation for both experimental and computational biomedical research, enabling discoveries spanning disease biology to precision medicine applications. However, as these banks scale rapidly, they face a fundamental bottleneck: Quality Control of physical and molecular integrity of samples [1]. Currently, physical integrity is assessed manually by pathologists a non-scalable approach that is subjective, time-consuming, and prone to inter-observer variability [2]; Molecular integrity is assessed using sequencing, costing both sample and finances, and again not scalable [2]. This bottleneck creates substantial barriers for establishing new biobanks with adequate quality assurance and limits the utilization of existing large biobank. While prior methods exist to address poor focus and staining imaging quality at scale [3], no scalable methods exist for automated assessment of biospecimen physical and molecular integrity quality metrics that determine research utility.

We present a computational pathology approach to determine both physical integrity (autolysis score) and molecular integrity (RNA Integrity Number, RIN score) from high-resolution histopathology images generated routinely during biobank development. Trained on publicly available GTEx cohort [4] comprising more than 30 million patch images from >25,000 non-diseased biopsies, we developed PathQC (Pathology-based Quality Control), a scalable way to quantify autolysis and RIN scores across 29 tissues. Our precise contributions are:

1. We first demonstrated that morphology in H&E encodes information for both RIN and autolysis scores. Here, we used recent advancements in digital pathology foundation models to extract morphological features. For 29 tissue types, we showed that morphology can predict autolysis and RIN scores (predicted vs. true correlation = 0.16-0.84 across tissues).
2. We provide a pan-tissue and easy-to-use PathQC GitHub package for community usage, where no information of tissue type is needed. Here, the input will be their slide and output will be quality control metrics.

This work addresses a critical infrastructure need in biomedical research. By enabling automated quality assessment of the millions of specimens in existing biobanks and facilitating quality-assured collection of new samples, we envision PathQC usage for both existing biobank quality assessment and accelerating the setting up of new ones. We thus envision that *npj Digital Medicine* audience will find this advancement of interest.

Sincerely,

Sanju Sinha, PhD

Assistant Professor

Sanford Burnham Prebys Medical Discovery Institute
